# A Bayesian framework for the analysis of systems biology models of the brain

**DOI:** 10.1101/468793

**Authors:** Joshua Russell-Buckland, Chris P. Barnes, Ilias Tachtsidis

## Abstract

Systems biology models are used to understand complex biological and physiological systems. Interpretation of these models is an important part of developing this understanding. These models are often fit to experimental data in order to understand how the system has produced various phenomena or behaviour that are seen in the data. In this paper, we have outlined a framework that can be used to perform Bayesian analysis of complex systems biology models. In particular, we have focussed on analysing a systems biology of the brain using both simulated and measured data. By using a combination of sensitivity analysis and approximate Bayesian computation, we have shown that it is possible to obtain a more complete understanding of the parameter space as compared to a maximum likelihood estimate based approach. This is done through analysis of simulated and experimental data. NIRS measurements were simulated using the same simulated systemic input data for the model in a ‘healthy’ and ‘impaired’ state. By analysing both of these datasets, we show that different parameter spaces can be distinguished and compared between different physiological states or conditions. Finally, we analyse experimental data using the new Bayesian framework and the previous maximum likelihood estimate approach, showing that the Bayesian approach provides a more complete understanding of the parameter space.

**Author summary:** *Systems biology* models are mathematical representations of biological processes that reproduce the overall behaviour of a biological system. They are comprised by a number of parameters representing biological information. We use them to understand the behaviour of biological systems, such as the brain. We do this by fitting the model’s parameter to observed or simulated data; and by looking at how these values change during the fitting process we investigate the mechanisms controlling the behaviour of our system. We are interested in understanding differences between a healthy and an injured brain. Here we outline a statistical framework that uses a *Bayesian* approach during the fitting process that can provide us with a distribution of parameters rather than single parameter number. We apply this method when simulating the physiological responses between a healthy and a vascular compromised brain to a drop in oxygenation. We then use experimental data that demonstrates the healthy brain response to an increase in arterial CO2 and fit our brain model predictions to the measurements. In both instances we show that our approach provides more information about the overlap between healthy and unhealthy brain states than a fitting process that provides a single value parameter estimate.

## Introduction

Systems biology models are used to understand complex biological and physiological systems comprised of large numbers of individual elements that give rise to emergent behaviours. These complex systems are dependent on both the properties of the whole network and on the individual elements [1]. This inherent complexity within the models can lead to difficulties in determining how best to interpret information obtained through their use.

At University College London, the family of BrainSignals models (and the BRAINCIRC model on which they are based) are used to understand the brain’s dynamics via a systems biology approach. They bring together a number of mathematical models relating to different aspects of blood circulation, oxygen transport and oxygen metabolism within the brain in order to develop a more complete model that can be used alongside experimental data to simulate physiological phenomena of the brain, such as autoregulation and neural activation. This allows us to understand how our measurements are linked to specific brain physiological and metabolic mechanisms.

All of the models were developed to reproduce broadband near-infrared spectroscopy (NIRS) measurements of brain tissue concentration changes of haemoglobin (oxygenation and haemodynamics) and cytochrome-*c*-oxidase (mitochondrial metabolism) and vary in their complexity and scope. Table 1 compares the number of reactions, equations, relations, reactions, variables and parameters in three different models. The first model developed was the ‘BRAINCIRC’ model in 2005 [2]. This built on an earlier circulatory model by Ursino and Lodi [3] and combined models for the biophysics of the circulatory system, the brain metabolic biochemistry and the function of vascular smooth muscle. This model was succeeded by the ‘BrainSignals’ model [4], which simplified the previous ‘BRAINCIRC’ model and added a submodel of mitochondrial metabolism. A number of additional versions were then developed from this, such as the ‘BrainPiglet’ model [5] which was developed to to simulate the physiological and metabolic processes of the piglet brain often used as the neonatal preclinical model. It involved modifying the default values for 11 of the 107 parameters used and was extended to include simulated measurements for magnetic resonance spectroscopy values that included brain tissue lactate and ATP production, measurements of which are available in piglet studies. This was extended in BrainPiglet v2 to incorporate the effects of cell death during injury [6]. In 2015, Caldwell et al. modified the BrainSignals model to produce the ‘BrainSignals Revisited’ model [7]. This made various simplifications to the BrainSignals model in order to reduce complexity and decrease the time taken to run a simulation, whilst being able to reproduce the same results and behaviour of the original model. This reduced model of the adult brain was later extended to simulate extracerebral haemodynamics to investigate confounding factors with brain near-infrared spectroscopy measurements, the ‘BSX’ model [8]. These models are run using the Brain/Circulation Model Developer environment (BCMD) and are defined in a simple text language.

**Table 1.** Comparison of the number of reactions, equations, relations, reactions, variables and parameters in the BRAINCIRC, BrainSignals Revisited and BrainPiglet v2.0 models.

|  | BRAINCIRC | BrainSignals Revisited | BrainPiglet v2.0 |
| --- | --- | --- | --- |
| Reactions | 81 | 5 | 17 |
| Differential Equations | 5 | 9 | 21 |
| Algebraic Relations | 72 | 3 | 3 |
| Variables | 168 | 40 | 128 |
| Parameters | 697 | 139 | 227 |

The data collected and analysed with the models primarily consists of broadband NIRS data, providing information about tissue oxygenation, through monitoring of oxy- and deoxy-haemoglobin levels, and cellular metabolism, through the concentration of cytochrome-c-oxidase. This data is then supplemented by systemic information such as blood pressure, arterial oxygen saturation and/or partial pressure of CO_2_.

The models are driven with input signals, such as the blood pressure and/or oxygen saturation, and simulate brain tissue measurments of oxygenation, blood volume and metabolism, as well as the middle cerebral artery velocity (Vmca) and the cerebral metabolic rate of oxygen (CMRO_2_). The model can be split into roughly 3 compartments - blood flow, oxygen transport and metabolism - with boundaries chosen to minimise interdependence. Fig 1 outlines this in more detail. One of the main uses of the models is to fit the model simulations to clinical and experimental data and investigate how model parameters are affected. In the case where data is collected from an injured or sick patient, these changes may illuminate what the underlying causes/mechanisms are behind the illness or injury is. For example, Caldwell et al. used BrainPiglet v2.0 to investigate why two piglets showed different recoveries following hypoxia-ischaemia, finding that the differences could be explained by including cell death within the model [6].

**Fig 1.**
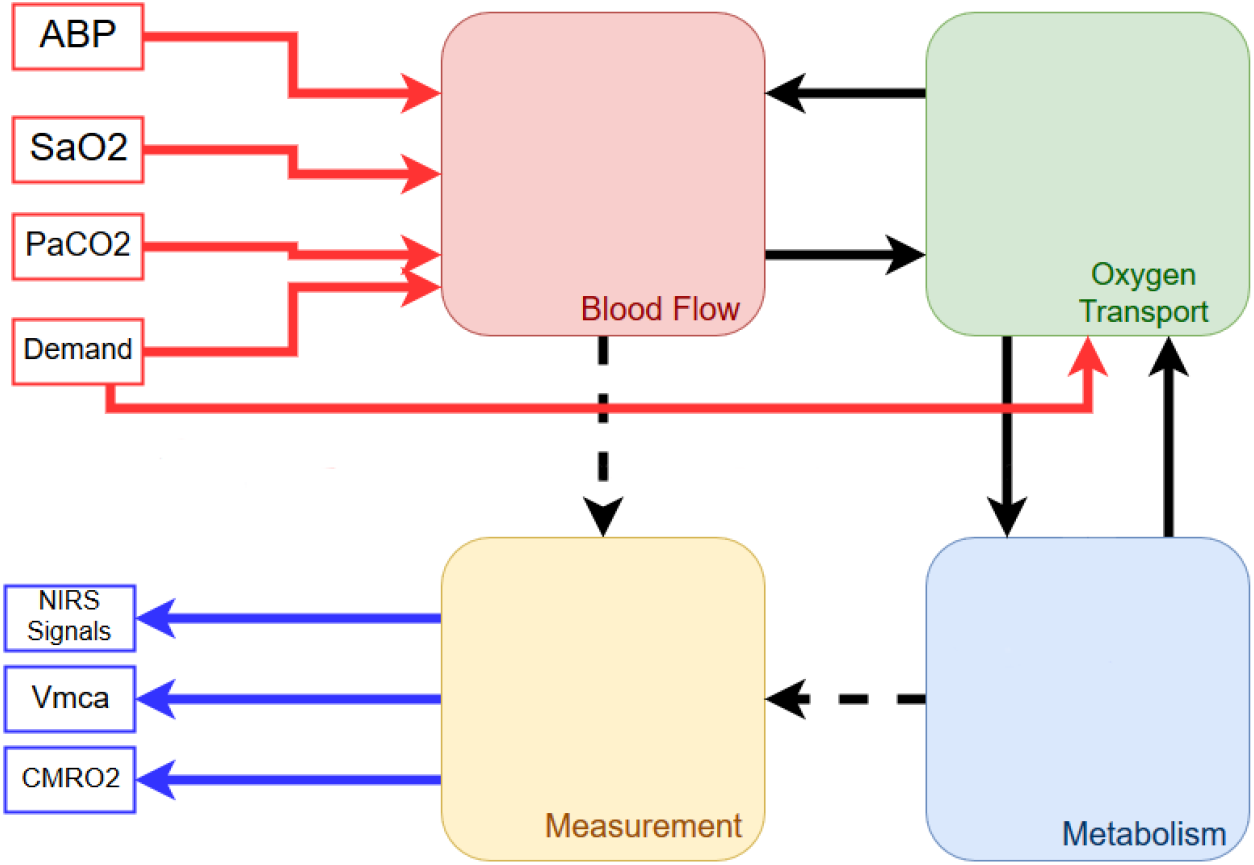
Simplified structure of a typical BrainSignals model. A typical BrainSignals model can be split into four compartments or submodels. The *blood flow* submodel represents blood flow from arteries to veins via the capillary bed and the *oxygen transport* submodel estimates diffusion of dissolved O_2_ from the capillary blood to the brain tissue. Delivered oxygen is then utilised by the *metabolism* submodel. Finally, the *measurement* submodel translates the internal states of the blood flow and metabolism submodels into observable outputs. Model inputs are shown in red and consist of arterial blood pressure (ABP), arterial oxygen saturation (SaO2), partial pressure of CO_2_ (PaCO2) and a parameter specifying relative demand, whilst measurable outputs are shown in blue, including NIRS signals as well as middle cerebral artery velocity (Vmca) and cerebral metabolic rate of oxygen (CMRO_2_).

The models are currently fit using a maximum likelihood based method, with a single value obtained for each parameter. Sensitivity analysis performed on the models to determine which parameters are most important in influencing each model output for any particular dataset. These parameters are then optimised using the PSwarm method [9] to minimise a given error metric, such as the Euclidean distance, between the modelled and measured signals. Through this each output has a set of optimised parameter values. Parameter values were limited to the same ranges used in the sensitivity analysis [6].

This approach has a number of drawbacks. The models are mechanistic and, if fitted to single value parameter estimates, will produce the same output for the same input. Physiology and biology, however, is unlikely to operate in such a constrained manner. In effect, the maximum likelihood approach, which produce single value estimates of parameters, can lead to overfitting of the model to the data. Additionally, this set of best-fit parameters for the model may not be representative of the full parameter space [10]. In an attempt to try and compensate for this potential drawback, Caldwell et al. [6] fit the BrainPiglet model multiple times for two different piglets and found that, whilst parameter values can vary within the same data, separate parameter spaces for each piglet did seem to exist based on the brain physiological status of the piglet following a hypoxic-ischaemic insult.

One of the key ways in which these models are used to extract information from data is through the use of parameter estimation and fitting. However, this step remains a difficult mathematical and computational problem, potentially originating in the lack of identifiability [11]. In addition, there has been discussion of ‘universal sloppiness’ within dynamic systems biology models. Gutenkunst et al. [12] proposed that *sloppiness*, where the parameters of a dynamic model can vary by orders of magnitude without affecting model output, is a universal property of systems biology models. Due to this sloppiness, it may not be possible to make parameter estimations that can be used to make inferences about the system [12, 13]. Chis et al. have stated however that sloppiness is not equivalent to a lack of identifiability and that a sloppy model can still be identifiable [14]. Apgar et al. note that experimental design can be used to constrain a sloppy parameter space by choosing a set of complementary experiments [15].

The use of a Bayesian methodology, by avoiding point estimates, can allow the full uncertainty of the problem to be captured [10]. In fact, the use of an Approximate Bayesian Computation (ABC) approach, discussed below, is particularly well suited to these kinds of problems [16]. There are many examples of Bayesian methods being used to analyse bioinformatics data and systems biology models [17], including in sequence analysis [18], gene microarray data [19] and in models of genetic oscillators [20] and DNA network dynamics [21]. There are a number of models that take a systems biology approach towards understanding physiology, particularly oxygen transport and blood flow, including the previously mentioned BrainSignals [2, 4, 6] and BrainPiglet [5, 6] models, the Aubert-Costalat model [22], and work by Fantini [23–26] and Orlowski and Payne [27, 28] where Bayesian parameter estimation could also be applied but has yet to be.

Bayesian inference utilises Bayes’ rule,

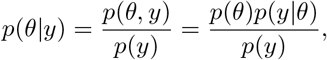

where *p*(*y*) = *∫_θ_ p*(*θ*)*p*(*y|θ*) *dθ* is the marginal probability of *y* and *p*(*y|θ*) is the *likelihood*. Typically, *p*(*y*) is not known and the likelihood will not be known explicitly or may require marginalising over some values of *θ*. This often leaves the solution analytically intractable. Instead we can try solve for *p*(*θ|y*) using a Monte Carlo or Markov Chain Monte Carlo (MCMC) approach.

Where a likelihood function can be defined there are a number of these methods that can be used to infer a posterior distribution, *p*(*θ|y*). The simplest is the Gibbs Sampler [29], which in its most basic form is a special case of the Metropolis-Hastings algorithm [30]. If a likelihood expression is unobtainable, as is the case with the BrainSignals models, a likelihood-free approach using ABC is required instead. There are a number of different methods available with the simplest being the ABC rejection algorithm (ABC REJ) approach.

The aim of this paper is to introduce the new *BayesCMD* modelling platform that can be used in systems biology models of physiology such as the BrainSignals models, but that can be replicated beyond these. For this work, we have chosen to use ABC REJ as whilst it is less efficient than the other methods mentioned here, the simplicity with which it can be implemented is a significant factor. The models and modelling environment used are already complex and so this initial work focuses on the use of the simplest method as proof of utility. We will demonstrate the effectiveness of this approach by using it to analyse two simulated datasets chosen to represent healthy and impaired brain states, before then using it on experimental data from a healthy subject undergoing a hypoxia challenge. We will show that the Bayesian approach allows us to extract more information from our data than the previous maximum likelihood approach, with a more complete picture of the parameter space being obtained.

## Materials and methods

Figure 2 shows a generalised outline of the final Bayesian analysis process. It can be split into three main sections: sensitivity analysis, Bayesian analysis and model checking. However, before applying the process, data must be generated or collected and an appropriate model chosen.

**Fig 2.**
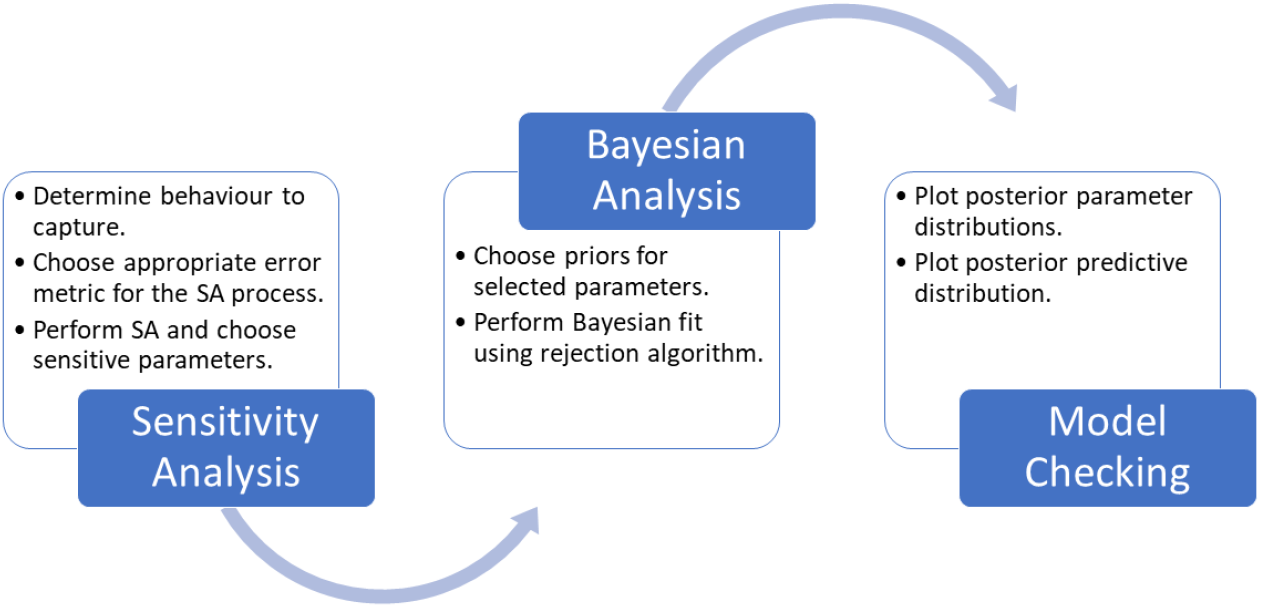
Generalised analysis process. A simplified representation of the Bayesian analysis process.

### Choice of Model

In this work we have chosen to use the refactored BrainSignals model [7], with a minor modification to include the haemoglobin difference (ΔHbO_2_ − ΔHHb = ΔHbD) as a model output alongside the normal outputs of oxyhaemoglobin (ΔHbO_2_), deoxyhaemoglobin (ΔHHb), total haemoglobin (ΔHbO_2_ + ΔHHb = ΔHbT), tissue oxygenation index (TOI), and cytochrome-*c*-oxidase (ΔCCO). Both ΔHbD and ΔHbT are included in the experimental dataset due to them being good indicators of brain oxygenation changes and brain blood volume changes respectively, with both being easily measured using broadband NIRS. All NIRS outputs, except TOI, are measured as changes relative to an initial value and therefore both data and model outputs are normalised to an initial value of 0.

### Data

Three datasets were used to test the new Bayesian model analysis process. Firstly, ‘healthy’ data was simulated using the BrainSignals model with the default parameter settings, as per [2, 4]. Next, the same inputs were used but with the model modified to represent an ‘impaired’ brain. To do this, a single parameter was changed to reflect a potential pathology or injury, to generate an ‘impaired’ simulated dataset. Finally, we used experimental data from a healthy adult undergoing a hypoxia challenge.

#### Simulated Data

Partial pressure of CO_2_ (PaCO_2_) and arterial blood pressure (ABP) were kept at their baseline values of 40 mmHg and 100 mmHg respectively, whilst arterial oxygen saturation (SaO_2_) was varied to simulate hypoxia through a decrease in arterial oxygen saturation from 97% to 65%. Initially, all model parameters were kept at their default values in order to simulate a healthy brain’s response to this challenge. Figure 3 shows the arterial saturation data and the model response across all considered model outputs.

**Fig 3.**
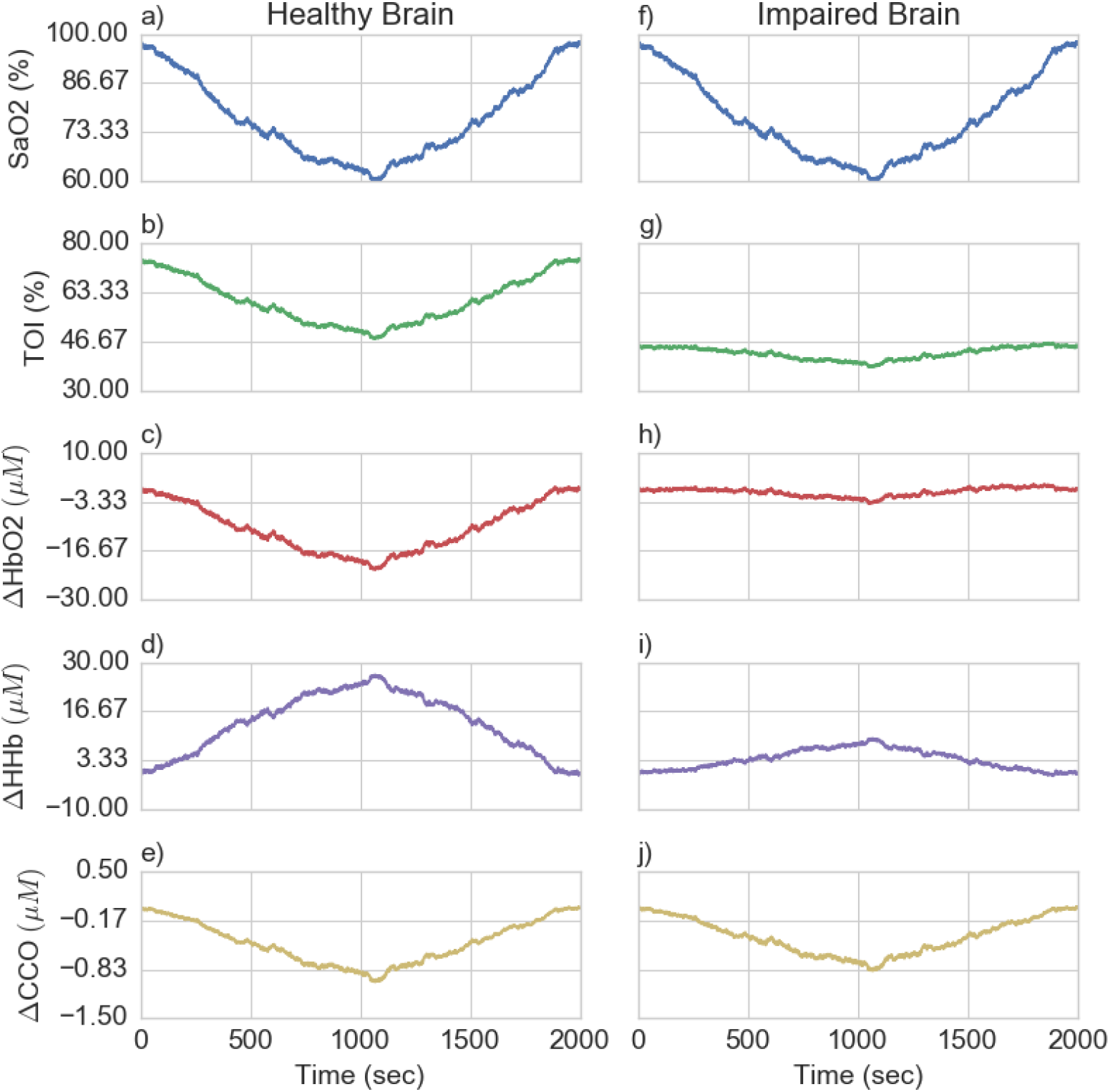
Healthy and impaired brain simulations. Figures a)-e) show simulations of a healthy brain’s response to hypoxia, whilst f)-j) show the impaired brain’s response. The input variable of arterial oxygen saturation is shown in blue and is the same for both simulations, whilst the outputs of TOI, ΔHbO_2_, ΔHHb and ΔCCO clearly differ between the two brain states.

After simulating the healthy brain response and determining its posterior parameter distribution, the model was altered to include a pathological or impaired brain state. This was done by changing a single parameter to be outside of the healthy parameter space. `r_t`, which affects the shape of the muscular tension relationship, was found to be sensitive in both the sensitivity analysis process (see Section and the Bayesian analysis. This is clearly seen in its comparatively narrow marginal posterior for the healthy data. Stiffening of blood vessels in the brain has also been noted as a potentially important factor in a number of different pathologies, including Alzheimers [31], and in autoregulation, as seen in 4.

The muscular tension relationship is defined as

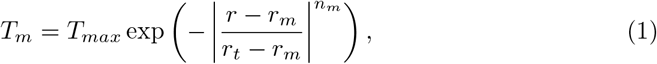

where *T_m_* is the muscular tension within the vessel wall and has a bell-shaped dependence on the vessel radius, taking value *T_max_* at some optimum radius *r_m_*. *r_t_* and *n_m_* are parameters determining the shape of the curve. Figure 4a illustrates the effect of changing *r_t_* on the shape of the curve and shows that decreasing *r_t_* leads to increased muscular tension for the same vessel radius due to a widening of the bell-shaped curve. This can be seen to represent a stiffening of vessels within the brain.

**Fig 4.**
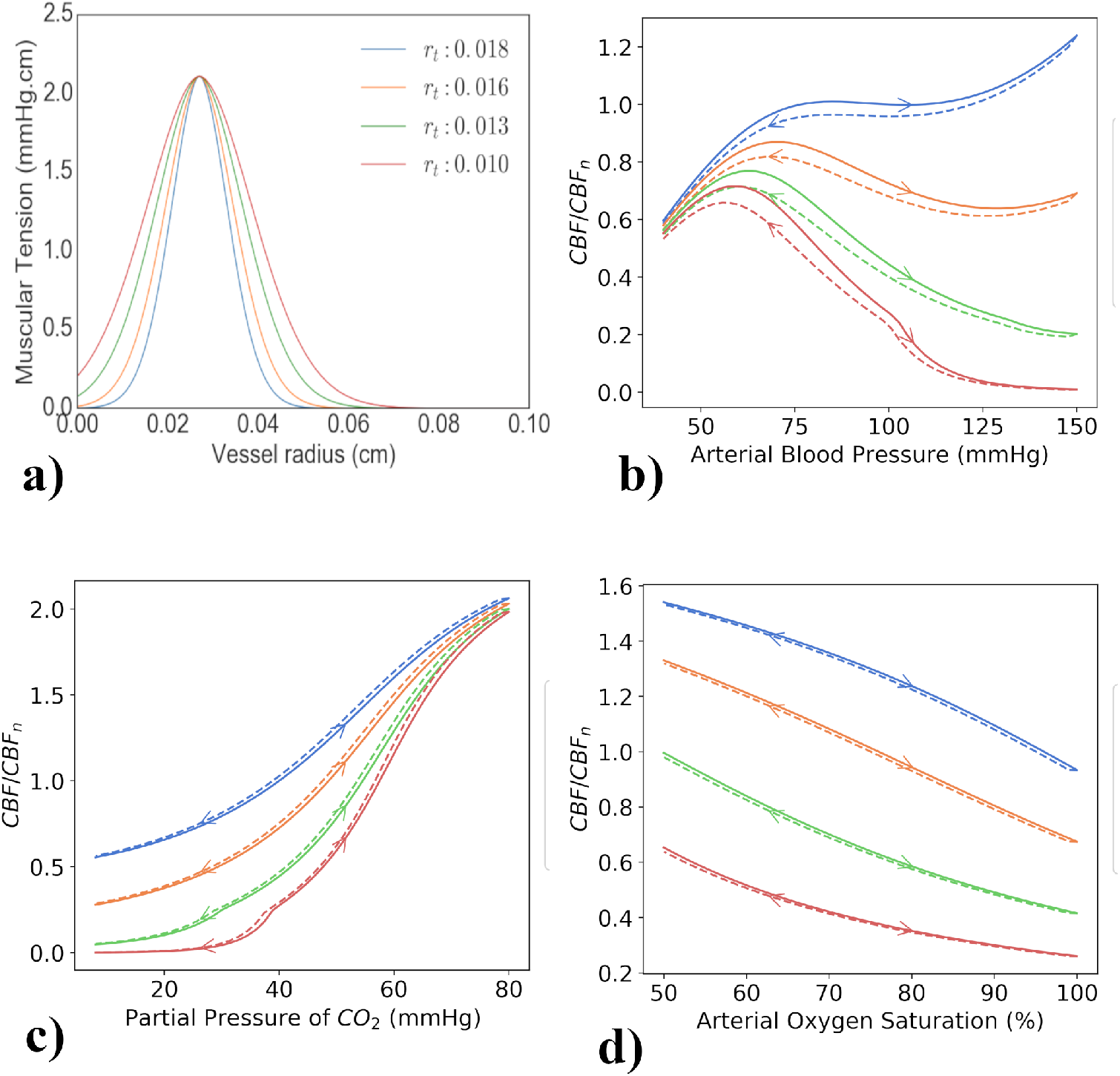
Figure 4a shows the effect of different *r_t_* values on the shape of the muscular tension curve for a range of vessel radii. It can be seen that reducing *r_t_* widens the curve, leading to increased muscular tension for the same vessel radius. Figures 4b, 4c and 4d show the effect of both increasing and decreasing model inputs on cerebral blood flow for different values of *r_t_*. Cerebral blood flow (CBF) is given as a proportion of the normal CBF (40 ml 100g^−1^ min^−1^). Changing *r_t_* has a significant effect on the brain’s ability to autoregulate within the model. Figure 4b shows that higher blood pressures causes a decrease in cerebral blood flow for lower *r_t_*, as opposed to an increase at the normal value of *r_t_* = 0.018 cm. Figure 4c shows that for lower *r_t_* values, CBF decreases quicker as PaCO_2_ is decreased. Figure 4d shows that across all considered oxygen saturations, lower *r_t_* gives a lower CBF.

Changing *r_t_* has a significant effect on the brain’s ability to autoregulate within the model as seen in figures 4b, 4c and 4d. Figure 4b shows that higher blood pressure causes a decrease in cerebral blood flow (CBF) for lower *r_t_* values, as opposed to an increase at the normal value of *r_t_* = 0.018 cm. Figure 4c shows that CBF is lower and decreases quicker for lower *r_t_* values as PaCO_2_ is decreased and figure 4d shows that across all considered oxygen saturations, lower *r_t_* gives a lower CBF.

Figures 3f)-j) shows the model response across all considered model outputs for this impaired brain state. The response of the model outputs to the same change in arterial saturation is much smaller than in the healthy simulation, with the TOI having a lower baseline value of around 45% as compared to around 75%.

#### Experimental Data

Experimental data will inherently contain more uncertainty for parameter fitting than data generated by the model itself. This makes it important to test the Bayesian analysis process on experimental data as well as that simulated from the model. The data used was originally collected by Tisdall et al. [32] and is shown in figure 5. Healthy adult humans had their arterial oxygen saturation reduced from baseline to 80%, whilst minimising changes in end tidal carbon dioxide tension (EtCO_2_).

**Fig 5.**
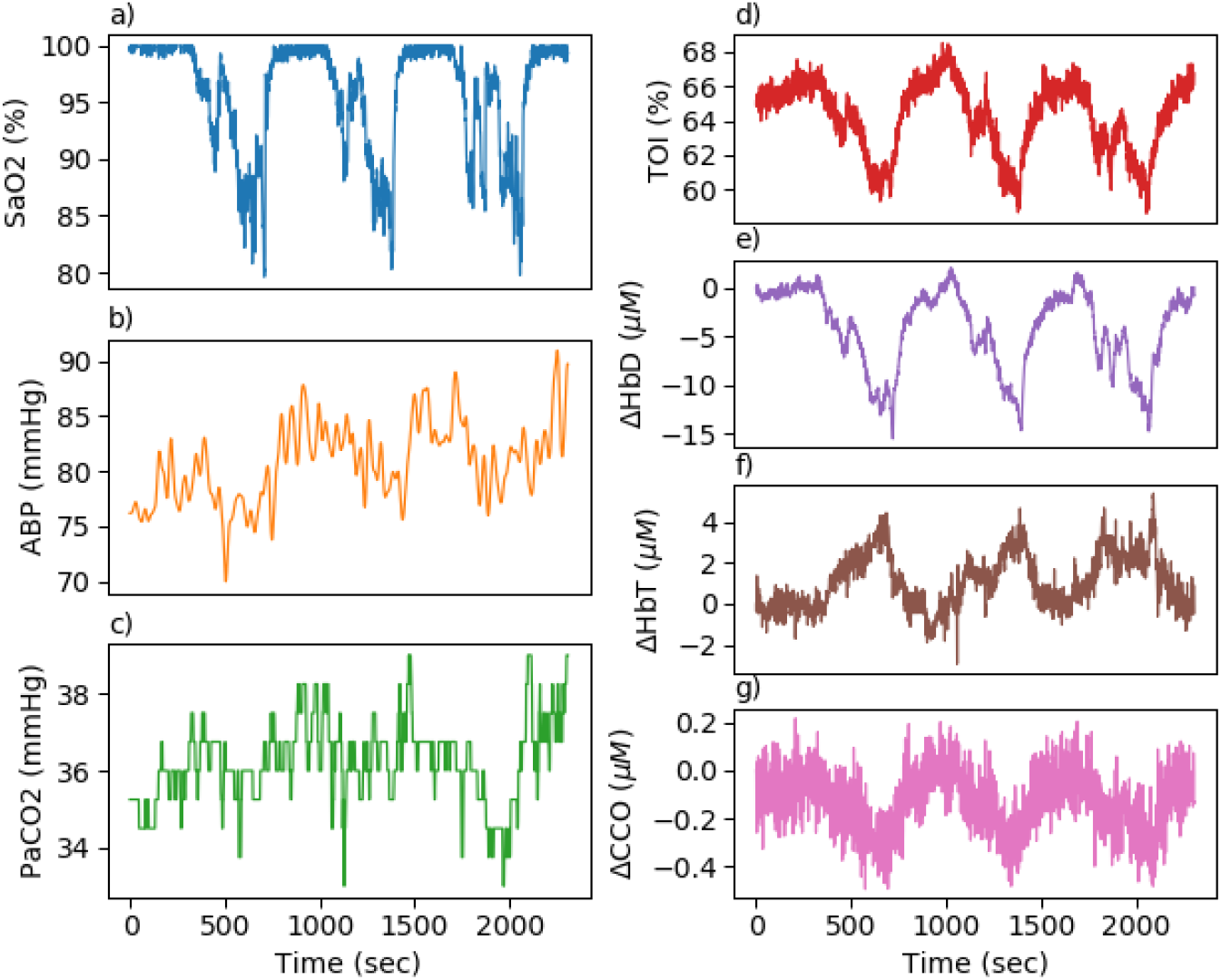
Experimental hypoxia data. Data collected from a healthy adult during a hypoxia challenge. Systemic data used as model inputs are shown in figures a), b) and c), with broadband NIRS measurements shown in figures d), e), f) and g).

The dataset contains three model inputs: arterial oxygen saturation, end tidal CO_2_ and arterial blood pressure, with EtCO_2_ converted to partial pressure of CO_2_. Blood pressure data was filtered using a low pass 5th order Butterworth filter, with a cut off of 0.05 Hz, to remove noise.

In terms of model outputs, only NIRS signals were used: ΔHbD, ΔHbT, ΔCCO and TOI. All data was resampled to 1 Hz.

### Sensitivity Analysis

When fitting a model as complex as BrainSignals, it is important to reduce the number of parameters that are required to be fit. We expect that not all parameters will have a significant impact on the model output for given set of input data. Instead, we can attempt to reduce the number of considered parameters through sensitivity analysis. We used the Morris method [33, 34], which is known to work well with a large number of parameters. The method requires the time series to be reduced to a single number and identifies the parameters that have produce the most variance in this summary value. Previously, we have used the Euclidean distance over the whole time series as our summary value but this has a number of significant drawbacks.

If the summary measure is the distance across the whole time series, we’re failing to capture specific changes that we know to be physiologically important. In the case of our hypoxia simulation, for example, we want to select parameters that are important in controlling the overall change from baseline. Taking the Euclidean distance over the time series as a whole however does not prioritise this behaviour. Figure 6a shows data generated from the same toy model function

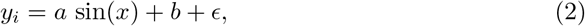

where *a, b* are both model parameters and *ϵ* is random Gaussian noise.

**Fig 6.**
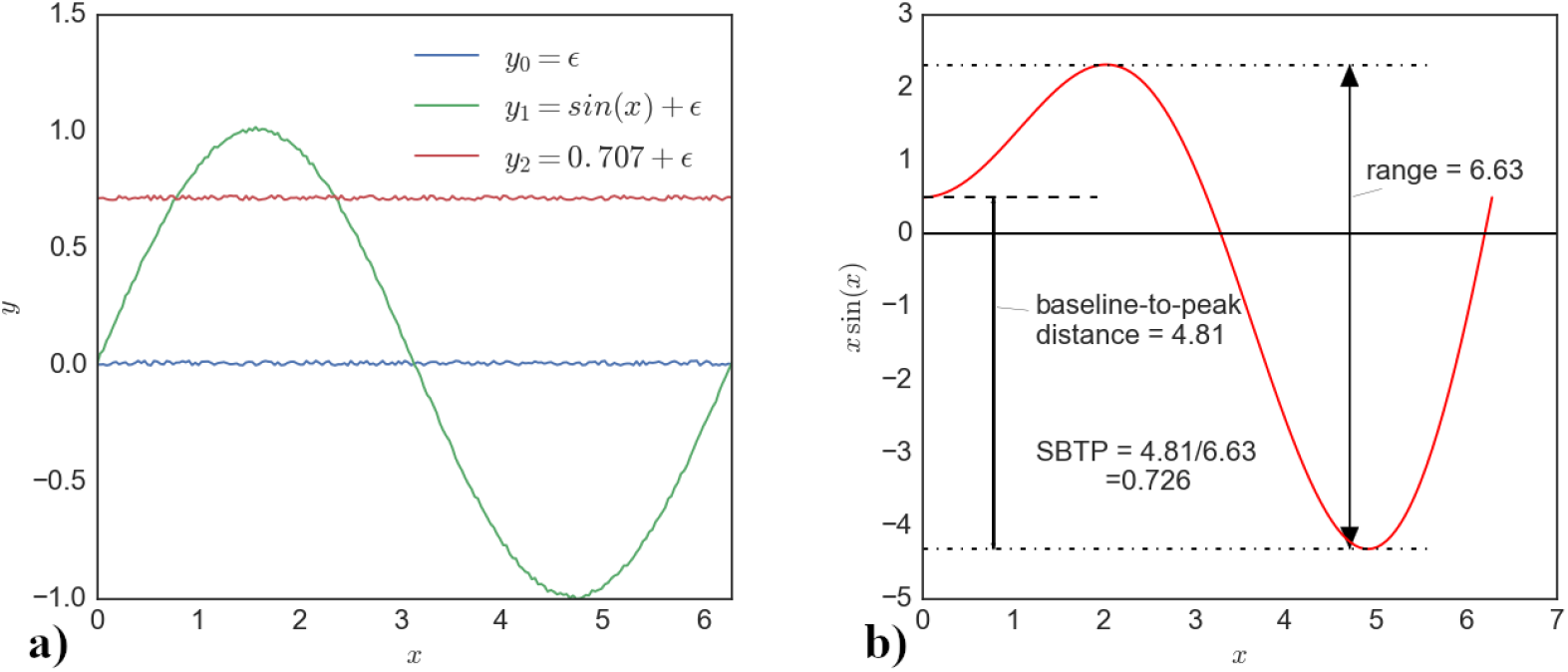
Figure 6a shows data generated from the same test function *y_i_* = *a* sin(*x*) + *b* + *ϵ*, where *a, b* are both model parameters and *ϵ* is random Gaussian noise. *x* was varied from 0 to 2*π*, producing data *y*_0_, *y*_1_ and *y*_2_ for the parameter sets Θ_0_: *a* = 0, *b* = 0, Θ_1_: *a* = 1, *b* = 0 and Θ_2_: *a* = 0, *b* = 0.707 respectively. Despite both *y*_1_ and *y*_2_ being qualitatively very different they are very similar when summarised using only the Euclidean distance, with *y*_1_ having a Euclidean distance *ε*_euc,1_ = 10.01 and *y*_2_ having a Euclidean distance *ε*_euc,2_ = 10.03. If we instead look at the scaled baseline-to-peak (SBTP) distance we find that *y*_1_ has a SBTP distance *ε*_SBTP,1_ = 49.98 and *y*_2_ has a SBTP distance *ε*_SBTP,2_ = 0.09. Figure 6b illustrates how the scaled baseline-to-peak distance is defined. The baseline-to-peak distance is the absolute distance from the baseline to max ({|*y_max_*|, |*y_min_*|}), This is then divided by the range of the data to get the distance as a proportion of the total change seen within the data. In this example, baseline-to-peak distance is 4.81 and the range is 6.63, giving a SBTP distance of 0.726.

Assume that we are trying to identify the parameter that is most important for determining how sinusoidal our model output is because we want to fit our model to this data. We decide to undertake sensitivity analysis, using a distance measure of some kind as our summary statistic that we can then pass to the sensitivity analysis method and the parameter found to be most sensitive will be the one we fit.

To generate our data *x* was varied from 0 to 2*π*, producing data *y*_0_, *y*_1_ and *y*_2_ for the parameter sets Θ_0_: *a* = 0, *b* = 0, Θ_1_: *a* = 1, *b* = 0 and Θ_2_: *a* = 0, *b* = 0.707 respectively, with output *y*_0_ and parameters *x*_0_ being our baseline. This is seen in figure 6a. It is clear from the figure that the two outputs *y*_1_ and *y*_2_ show very different behaviour, with *y*_1_ being the behaviour we want to optimise for.

Despite both *y*_1_ and *y*_2_ being qualitatively very different they are very similar when summarised using only the Euclidean distance, with *y*_1_ having a Euclidean distance *ε*_euc,1_ = 10.01 and *y*_2_ having a Euclidean distance *ε*_euc,2_ = 10.03.

Instead we can define a new summary measure, which we will call the “scaled baseline-to-peak” (SBTP) distance. We know that we want to find the parameter that determines how sinusoidal our model is. One way to emphasise this behaviour is to find the distance from our baseline to the maximum or minimum (whichever has the largest absolute value) of our data, as illustrated in Figure 6b. We then scale this by the range of our data to normalise it and avoid issues comparing data of different magnitudes. This gives us

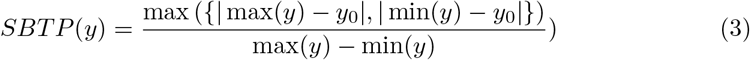

We then find the Euclidean distance between *SBTP* (*y*_1_) and *SBTP* (*y*_2_)

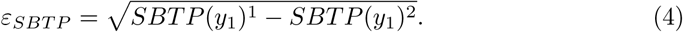

If we use this Euclidean distance between scaled baseline-to-peak (SBTP) distances, *ε_SBTP_*, we find that *y*_1_ has a distance *ε*_SBTP,1_ = 49.98 and *y*_2_ has a distance *ε*_SBTP,2_ = 0.09. Here we see that the difference in behaviour is clearly highlighted.

We scale our baseline-to-peak distance because a number of model outputs significantly vary over different scales. For example, cerebral oxygenation can be measured through TOI which is a percentage and, as seen in figure 3 can vary over 10-20%. Cytochrome-*c*-oxidase however, varies over a much smaller range, with a change of less than 1 µm being typical. Failing to account for these different scales will lead to parameters that affect larger magnitude outputs being identified as more sensitive than those that affect smaller magnitude outputs, even if the relative change is significant.

For example, if changing a parameter *θ*_1_ causes the CCO change seen in figure 3e) to double to a minimum of −2 µm, whilst a change in a parameter *θ*_2_ causes TOI to decrease to 55%, without scaling the model seems more sensitive to *θ*_2_ because the magnitude of the change is much more, even though the relative change is smaller. If we consider this change proportional to the range of our data however, we account for its relative size.

It should also be noted that this choice of metric is specific to the behaviour being optimised for. For example, in the case of a signal that is non-oscillatory, a different summary method would be required based around the behaviour to be replicated within that particular signal.

We used the Morris elementary effect method [33] variant devised by Saltelli et al. [35]. This provides us with two notable statistics: the mean of the absolute values of the changes, µ_*_, and their standard deviation, *σ*. The larger the value of µ_*_, the more influential parameter is on the output, whilst the larger the standard deviation, the more non-linear the influence of the parameter is. The top ten most sensitive parameters were chosen to fit the model. The parameter range considered for sensitivity is the default value ±50%. Sensitivities are calculated for each output as well as across all outputs jointly. This joint sensitivity is calculated by summing the SBTP value for each output and then determining variability in this total.

### Approximate Bayesian computation

After selecting the most important parameters, the model was fit using the rejection algorithm [36]. This is defined, as per [37], as:

1. Sample a candidate parameter vector *θ^*^* from the proposal distribution *p*(*θ*).
2. Simulate a dataset *y^rep^* from the model described by a conditional probability distribution *p*(*y|θ^*^*).
3. Compare the simulated dataset, *y^rep^*, to the experimental dataset, *y*, using a distance function, *d*, and tolerance, *ϵ*. If *d*(*y, y^rep^*) *≤ ϵ*, accept *θ^*^*. The tolerance *ϵ ≥* 0 is the desired level of agreement between *y* and *y^rep^*.

The output of the ABC algorithm used will be a sample from the distribution *p*(*θ|d*(*y, y^rep^*) ≤ *ϵ*). If *ϵ* is sufficiently small, then *p*(*θ|d*(*y, y^rep^*) ≤ *ϵ*) will be a good approximation for the posterior *p*(*θ|y*).

The choice of *d*(·, ·) is important, just as with the sensitivity analysis. Previously the Euclidean distance has been used to fit the model but, as in the case of the sensitivity analysis, this fails to account for outputs that vary over different magnitudes. Instead, we have chosen to include a number of other distance metrics including the root-mean-square error (RMSE) and the normalised root-mean-square error (NRMSE). These are defined as

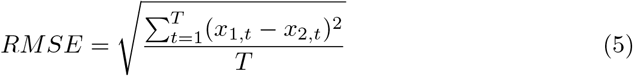

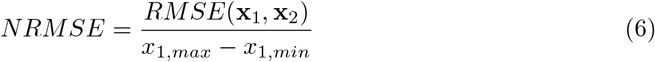

where **x**_1_ and **x**_2_ are the two time series being compared, running over *t* = 1 to *t* = *T*, with *T* being the total number of time points.

By dividing the RMSE by the range of the data, the errors for time series that vary over different magnitudes are comparable. Without doing this, parameters that mainly affect outputs that vary over larger magnitudes are preferentially optimised. Normalisation prevents overfitting of one output at the expense of others, providing a more reliable joint posterior distribution after fitting.

After an initial exploratory fitting of the different datasets, it was found that setting an absolute tolerance value was not a suitable selection criteria. This was due to massively differing distance values between datasets, with all parameter combinations in the simulated healthy dataset producing NRMSE values smaller than almost all parameter combinations on the impaired dataset.

In general, the number of accepted particles that gives an adequate approximation of the posterior distribution is problem dependent; dispersed posterior distributions will ultimately require more particles. Poor estimation of the posterior can in most cases result in a wide posterior predictive distribution which appears to give a poor quality fit because outlier posterior samples cause biases. To address this issue in a pragmatic way, a fixed acceptance rate of 0.01% was set. This meant the 0.01% parameter combinations with the lowest *d*(*y, y^rep^*) were used as the posterior. The posterior was visualised through kernel density estimation on a pairplot using the Seaborn plotting package [38]. The posterior predictive density is then generated by sampling directly from the posterior 25 times and the model simulated for each sample. The results are aggregated and plotted, with the median and 95% credible interval marked on the plot.

The model was run in batches of 10,000,000 and the parameter combinations within the acceptance rate were used as a posterior. This batch size was chosen as a compromise between sufficient sampling of the parameter space and the computational time required to run the batch. The quality of the fit obtained from this posterior determined if the model had been run a sufficient number of times to sample the posterior adequately. If the posterior predictive distribution failed to capture the behaviour seen in the “true” data, then the process was repeated until a more adequate fit was obtained.

## Results

### Sensitivity Analysis

#### Simulated Data

Sensitivity analysis was performed for the simulated healthy data set for the CCO, HbO_2_, HHb and TOI outputs. Figure 7 shows the sensitivity analysis results across all four outputs individually and for the outputs considered jointly. The results are plotted as bar charts, with sensitivity, as per the *µ_*_* value, on the x-axis.

**Fig 7.**
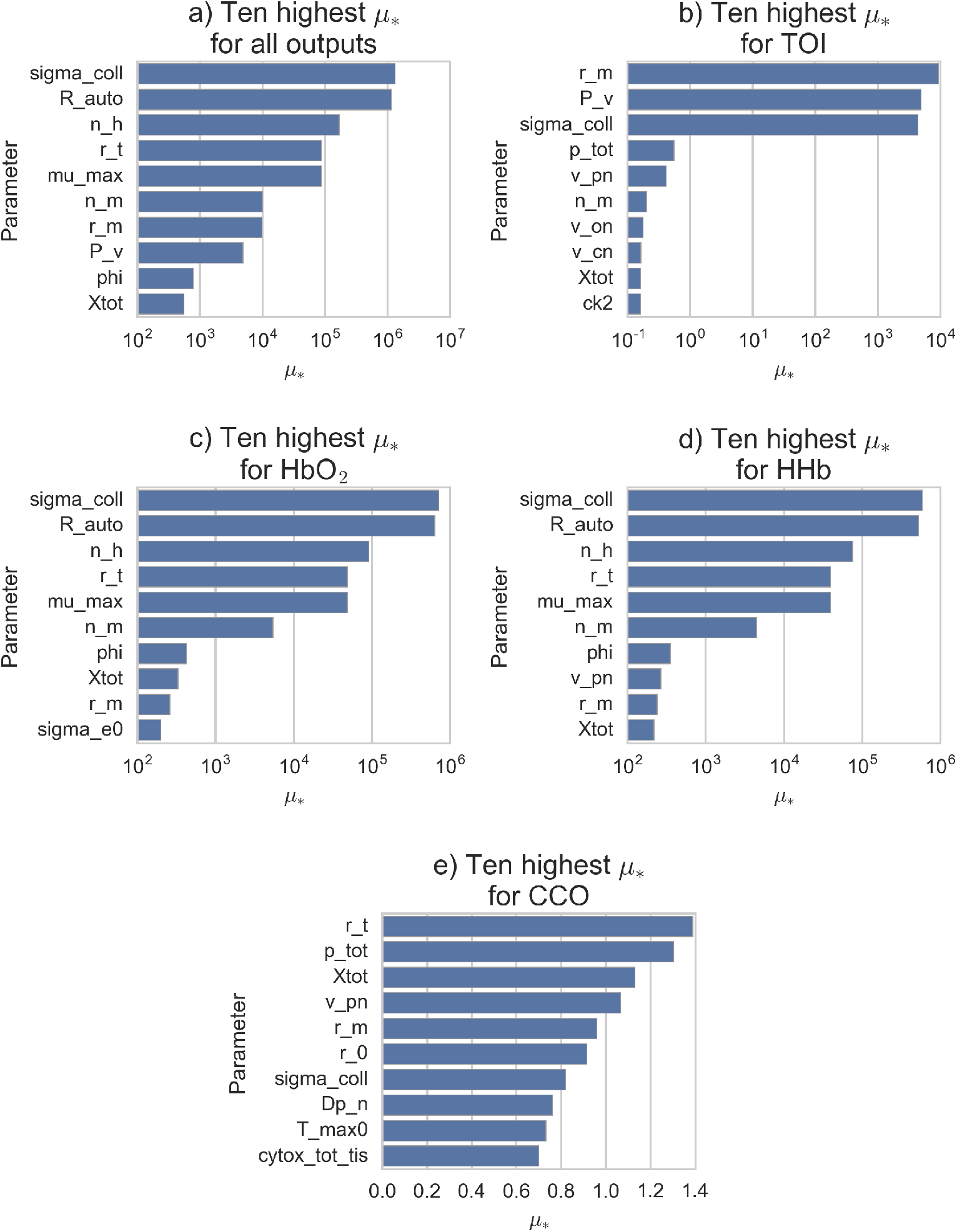
Sensitivity analysis across all outputs for simulated data set. Bar charts showing *µ_*_* for the 10 most sensitive parameters across all model outputs, with values plotted on a log scale where appropriate. Distance used for calculation is the sum of *ε_SBTP_* across all model outputs. All outputs except cytochrome-*c*-oxidase alone have *µ_*_* values that vary on a logarithmic scale.

Table 2 shows the selected parameters, their respective *µ_*_* values and their definitions and default values. The total sensitivity analysis results, shown in figure 7a, produced 10 parameters to be used in fitting the model. Sensitivity analysis based on individual outputs showed that different parameters were important for different outputs, with TOI, in figure 7b being dominated by `r_m, P_v` and `sigma_coll` and oxyhaemoglobin, in figure 7c, and deoxyhaemoglobin, in figure 7d, dominated by `sigma_coll` and `R_auto`. Cytochrome-*c*-oxidase however showed levels of dependence that were similar across many parameters, as seen in figure 7e, with *µ_*_* values falling within a range of 0.7. For all individual outputs and the combined output, only `Xtot, r_m` and `sigma_coll` were within the 10 most sensitive in all cases.

**Table 2.** Sensitivity analysis results for simulated data, including each selected parameter’s definition and default value. ^*^ See [7] and [4] for a full explanation of this parameter and the stimulus *µ*.

| Parameter | $\mu_*$ | Definition | Default value |
| --- | --- | --- | --- |
| sigma_coll | $1.32 \times 10^6$ | Pressure at which blood vessels collapse. | 62.79 mmHg |
| R_auto | $1.16 \times 10^6$ | Autoregulatory reactivity to oxygen. | 1.5 |
| n_h | $1.68 \times 10^5$ | Hill coefficient for oxygen dissociation from haemoglobin. | 2.5 |
| r_t | $8.82 \times 10^4$ | Radius in the muscular tension relationship. | 0.018 cm |
| mu_max | $8.82 \times 10^4$ | Upper bound for the transformed stimulus $\mu^*$. | 1 |
| n_m | $9.97 \times 10^3$ | Exponent in the muscular tension relationship. | 1.83 |
| r_m | $9.77 \times 10^3$ | Vessel radius at which muscular tension is maximal. | 0.027 cm |
| P_v | $4.88 \times 10^3$ | Venous blood pressure. | 4 mmHg |
| phi | $7.85 \times 10^2$ | Oxygen concentration at half-maximal saturation. | 0.036 mM |
| Xtot | $5.54 \times 10^2$ | Total concentration of haemoglobin O<sub>2</sub> binding sites in blood. | 9.1 mM |

#### Experimental Data

Sensitivity analysis was undertaken on the experimental dataset to determine the parameters to be fit. Table 3 shows the selected parameters, their respective *µ_*_* values and their definitions and default values. Figure 8 shows the results across all outputs. When considering all outputs jointly, the effect of `n_m` and `r_m` is significantly larger than all other parameters, but when looking at the individual outputs it’s clear that the other parameters are still important, but the magnitude of the impact `n_m` and `r_m` have on the overall variability is drastically larger.

**Table 3.** Sensitivity analysis results for experimental data, including each selected parameter’s definition and default value. *See [7] and [4] for a full explanation of this parameter and the stimulus *µ*. †This is the arterial PaCO_2_ input put through a first order filter to simulate varying time response and is typically the same as arterial PaCO_2_. For more information see [4]

| Parameter | $\mu_*$ | Definition | Default Value |
| --- | --- | --- | --- |
| n_m | $1.87 \times 10^7$ | Exponent in the muscular tension relationship. | 1.83 |
| r_m | $1.87 \times 10^7$ | Vessel radius at which muscular tension is maximal. | 0.027 cm |
| K_sigma | $1.01 \times 10^3$ | Parameter controlling the sensitivity of $\sigma_e$ to vessel radius $r_m$. | 10 |
| p_tot | $6.29 \times 10^3$ | Total protons removed from the mitochondrial matrix by the three modelled electron transport reactions. | 20 |
| k_aut | $3.65 \times 10^2$ | Parameter controlling overall functioning of autoregulatory response, with a value of 1 meaning intact autoregulation. | 1 |
| v_cn | $2.76 \times 10^2$ | Normal filtered $\text{PaCO}_2$†. | 40 mmHg |
| sigma_e0 | $2.37 \times 10^2$ | Parameter in the elastic tension relationship. | 0.1425 mmHg |
| k2_n | $2.17 \times 10^2$ | Normal forward reaction rate for the reduction of $a_3$. | $3915.68 \text{ s}^{-1}$ |
| Xtot | $1.48 \times 10^2$ | Total concentration of haemoglobin $\text{O}_2$ binding sites in blood (4 times the haemoglobin concentration). | 9.1 mM |
| R_autc | $1.31 \times 10^2$ | Autoregulatory reactivity to carbon dioxide. | 2.2 |

**Fig 8.**
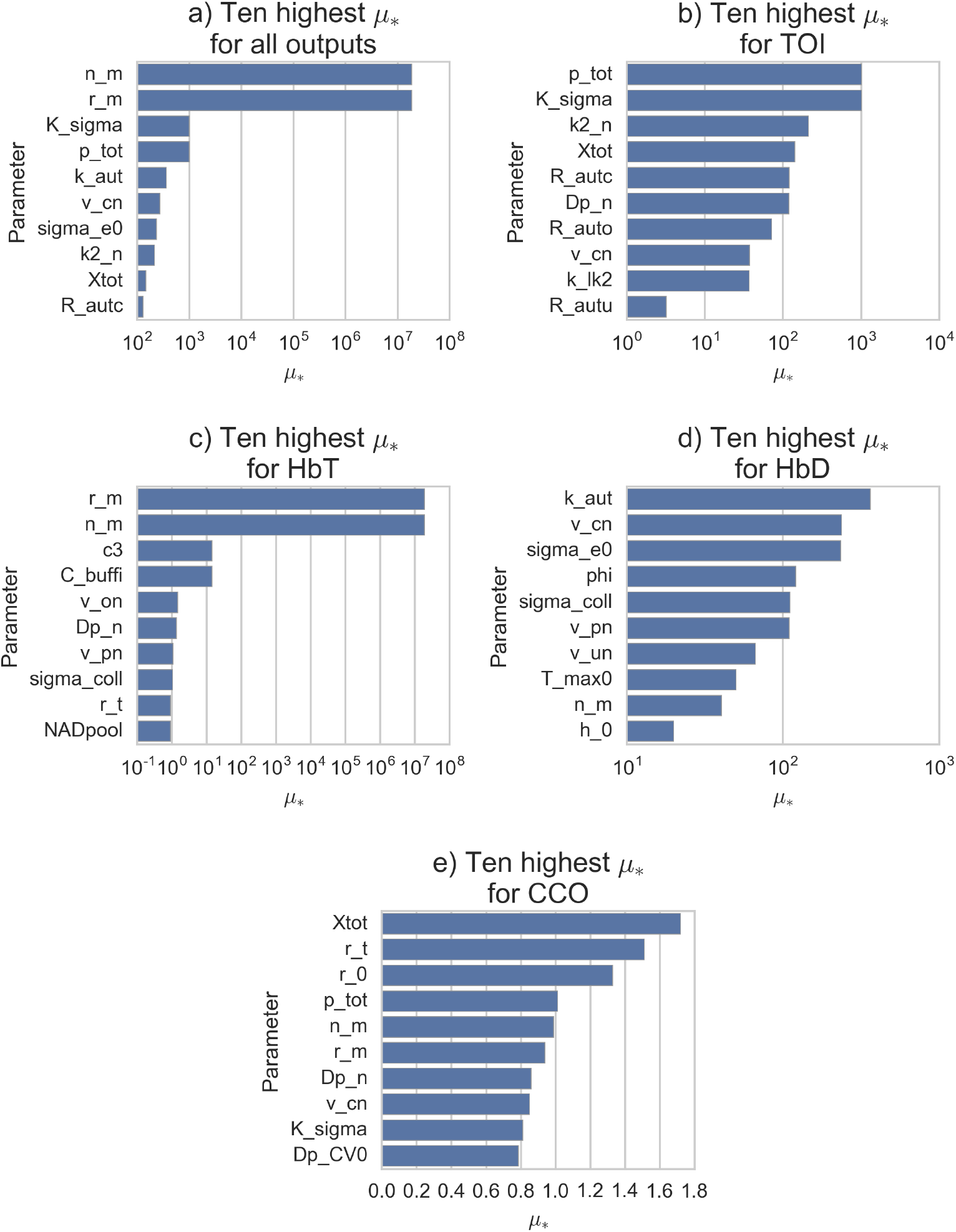
Sensitivity analysis across all outputs for experimental data set. Barplots showing *µ_*_* values for the 10 most sensitive parameters across all model outputs, with the x-axis plotted using a log scale where appropriate. Distance used for calculation is the sum of *ε_SBTP_* across all model outputs.

Unlike the simulated dataset, 9 of the top 10 most sensitive parameters have *µ_*_* values between approximately 10 and 1000 which is significantly smaller than the range of the *µ_*_* values for TOI in the simulated data.

Similarly, the most sensitive parameters for HbD fall within a very small range with no one parameter obviously determining the majority of the output’s behaviour. In contrast, the two most sensitive parameters for HbT, `r_m` and `n_m`, are approximately 10^6^ times larger than the third highest. As with the simulated data, *µ_*_* values for CCO have much smaller values than all other outputs and fall within a range of 1.0. Unlike the simulated data, no parameters were sensitive across all individual and joint outputs.

#### Parameters

Whilst a full exploration of the parameters within the BrainSignals model is outside the scope of this paper, we advise the reader to look at the original publications [4, 7], and provide a brief overview of some of those identified as important here.

A number of the parameters identified as being important for the above datasets, such as `R_auto` and `mu_max`, are dimensionless parameters. They are often model specific parameters that cannot be directly measured and instead need to be considered in the context of their meaning within the model. For example, an increase in `R_auto` would mean that the autoregulatory response would become more sensitive to changes in oxygen concentration. In contrast, other parameters such as `Xtot`, which is four times the concentration of haemoglobin, are more easily measured in an experimental or clinical setting.

Some of the parameters identified as important are linked closely to the shape of the autoregulatory response of the model and its sensitivity to changes in model inputs. These are `R_auto` and `mu_max` in the simulated dataset, and `v_on, v_un, R_autc` and `v_cn` in the experimental dataset. As we are driving the model with a changing input, the identification of these parameters as important seems physiologically sensible. It should also be noted that, despite other parameters not directly controlling the autoregulation response, the interconnected and complex nature of the BrainSignals model means that other parameters may still have an impact on it indirectly, for example the parameter `r_t` controls the stiffness of blood vessel walls, which is important in controlling blood flow during autoregulation.

More detailed information on the exact nature of these parameters and how they function within the BrainSignals model can be found in [7].

### Bayesian Analysis

#### Simulated Data

The BrainSignals model was fit to the simulated “healthy” dataset initially. The model was run 10,000,000 times before determining that the posterior had been estimated sufficiently well, based on the quality of the posterior predictive distribution. The particles in the posterior were found to have 0.019170 *≤ ε*_NRMSE_ *≤* 0.098098. Figure 9a shows this posterior distribution in blue. `Xtot, phi` and `r_t` show narrow marginal distributions whilst the others are much wider. Median values for all parameters are close to the model value, with `R_auto` showing a skew towards lower values in its marginal distribution that also leads to a median slightly lower than the model value. Figure 9b shows the posterior predictive distribution produced by sampling 25 times directly from the posterior, and shows a very good fit.

**Fig 9.**
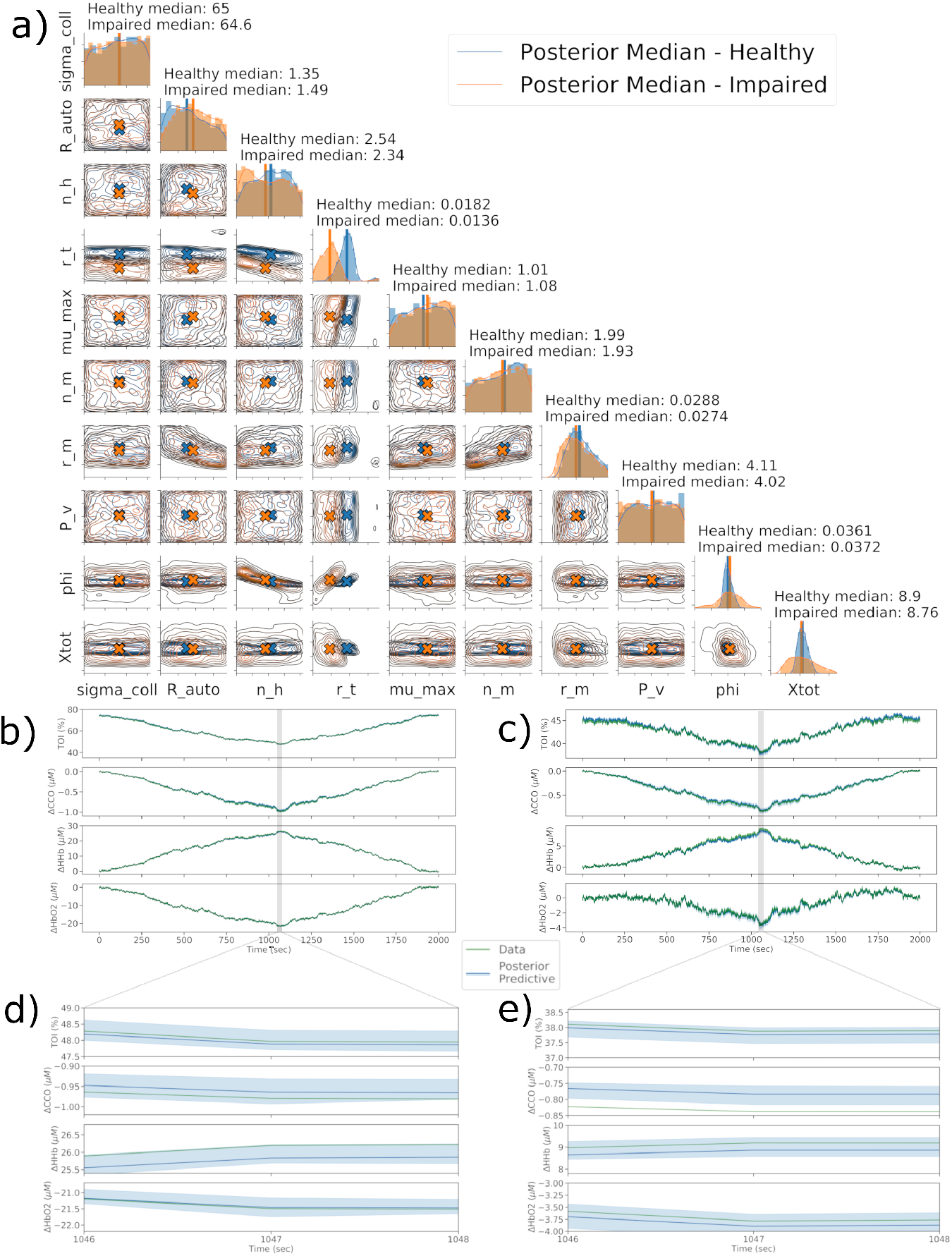
Comparison of errors, posteriors and predictions for healthy and impaired simulated data. Figure 9a shows the posteriors for healthy and impaired data based on an acceptance rate of 0.01%. Posterior are shown over the full prior range as defined in table 4 in S1 Table and table 5 in S2 Table. Figures 9b and 9c show the predicted time series data from the healthy and impaired posteriors respectively. Each posterior was sampled 25 times and the resulting runs aggregated, with the median and 95% credible intervals plotted in dark blue and light blue respectively. Figures 9d and 9e show a zoomed in view of each output in order to show the credible interval of the posterior predictive distribution.

This healthy posterior was then used to define an impaired brain, as mentioned above. `r_t` was set to 0.013 mmHg and the model driven with the same inputs as the healthy simulation. This “impaired” dataset was then fit using the same approach as above, using the sensitivity analysis results. The model was run 30,000,000 times, with the increased run number required in order to sufficiently estimate the posterior. With an acceptance rate of 0.01%, a posterior was produced based on 3000 particles having 0.019170 *≤ ε*_NRMSE_ *≤* 0.267152. Despite the higher error values as compared to the healthy data, the resulting fit was still deemed very good. Figure 9a shows this posterior in orange and figure 9c shows the time series generated by sampling 25 times directly from this posterior. `Xtot, phi` and `r_t` show marginal distributions that are narrower than the others, but wider than those seen in the healthy posterior. All parameters have median values close to the value set in the model. A separation between the healthy `r_t` and impaired `r_t` marginal distributions is clearly visible.

Figures 9d and 9e show a zoomed in view of each output in order to show the 95% credible interval of the posterior predictive distribution. This is not clearly visible on the full trace as it is reasonably small.

#### Experimental Data

When approaching the experimental data, the criteria for a good fit were different to those in the simulated dataset. With the simulated dataset, any parameters not chosen for fitting would have the same value during the fitting process as during the generation of the simulated dataset. In the experimental data however, it is almost certain that the default values of any parameters not chosen for fitting would not have the exact same value as their biological, real-world analogue. As a result, instead of looking for a perfect fit, we instead look for qualitative behaviours to be reproduced, such as the periodic increase and decrease in output values due to the repeated hypoxia challenges.

The fitting process required 20,000,000 runs before a satisfactory fit was obtained, and with an acceptance rate of 0.01% the posterior in figure 10a consisted of 2000 particles with 0.778492 *≤ ε*_NRMSE_ *≤* 0.802900. The model was also fit using the previous OpenOpt method, and the values obtained from that are also shown for comparison. We can see that for parameters with reasonably well defined posterior, the OpenOpt values and the posterior median are reasonably close, but for those showing a wider distribution, the OpenOpt value can vary massively from the posterior median. For `sigma_e0` and `k2_n` the OpenOpt value is at one extreme end of the prior range, whilst the median remains central due to the distribution being uniform. Figure 10b shows the predicted time series for all outputs based on the posterior shown in figure 10a. The posterior was sampled 25 times with the resulting time series aggregated, with the median and 95% credible intervals plotted. Overall behaviour is reflected in the predicted trace, with 3 distinct periods of hypoxia visible as periodic behaviour within all signals. Shown in green is the fit obtained using the OpenOpt method, which has an error *ε*_NRMSE_ = 0.77518. It is clear that both methods are able to achieve similar fits, but the Bayesian method provides more information about the space of possible parameter combinations and the resultant uncertainty in fitted model output.

**Fig 10.**
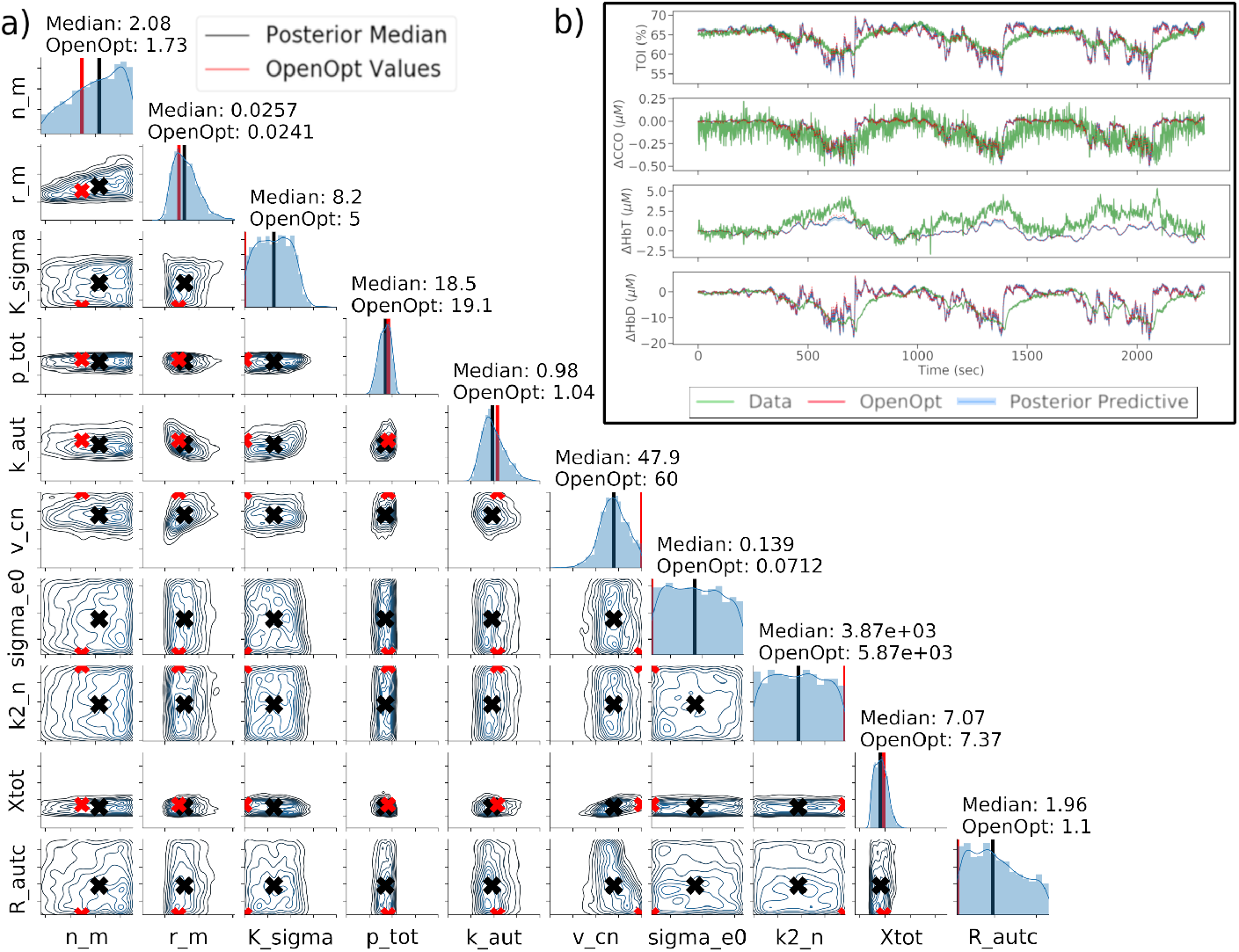
Errors, posterior and predicted fits for the experimental data set. Figure 10a shows the posterior distribution for the experimental data set, based on an acceptance rate of 0.01%. The posterior median is shown in black and the OpenOpt predicted value is shown in red. Posterior are shown over the full prior range as defined in table 6 in S3 Table Figure 10b shows the predicted time series for all output based on the posterior shown in figure 10a. The posterior was sampled 25 times with the resulting time series aggregated, with the median and 95% credible intervals plotted in dark and light blue respectively. Overall behaviour is reflected in the predicted trace, with 3 distinct periods of hypoxia visible as periodic behaviour within all signals. The fit obtained using OpenOpt is shown in red.

## Discussion

In this work we have introduced a new Bayesian analysis for interpretation of the BrainSignals models. The process was tested and used to analyse two simulated datasets and one experimental data set. The Bayesian approach provides us with complete information about the parameter space and takes into account the prior information we have about physiological parameters. Both of these factors are extremely important when drawing physiological conclusions from any parameter estimates.

We have shown how the method can be used to define healthy and impaired parameter spaces, as shown with the simulated datasets, and how for some parameters these spaces may overlap. We have also shown how the new Bayesian approach provides more information about the parameter space than the previous OpenOpt maximum likelihood method. Looking at only the healthy data set, the parameters `sigma_coll, P_v` and `mu_max` all have marginal posteriors with a median at the default value set in the model, but with distributions that cover the entirety of the prior distribution initially set. Determining that a parameter’s posterior distribution is not tightly constrained is important when drawing physiological conclusions from the model fitting process.

This is seen more clearly when looking at the experimental data. Many of the parameters show relatively narrow marginal posteriors, but `sigma_e0` and `k2_n`, which were both identified as important by the original sensitivity analysis, are both shown to have wide distributions, suggesting insensitivity within the prior range. The previous OpenOpt method produces an almost identical fit as the Bayesian approach but provides significantly less information about the parameter space. For `sigma_e0, k2_n, v_cn, R_autc` and `k_sigma` the OpenOpt values fall outside of the interquartile range of the posterior distribution, yet produce equivalent model simulations. If considering the OpenOpt estimate alone, it would be simple to draw the conclusion that these parameters have shifted away from the default ‘healthy’ value, showing some sort of physiological change during the hypoxia challenge. However, when we look at the posterior obtained through the Bayesian method, the median value is close to the default value and in fact parameter values across the entire prior range produce similar results. As a result we can instead say that for this data the model is insensitive to these parameters, with a median value that would be considered ‘healthy’.

There are a number of other methods that can also be used to explore and define the parameter space. The previously used maximum likelihood based method, for example, can provide estimates and confidence limits of parameter values, but under the assumption that the maximum likelihood estimator is normally distributed around the maximum. It may also be possible to use a profile likelihood [39], but whilst this will provide information about the distribution of the parameter space without assuming normality, it is computationally expensive and does not take into account prior information about the parameters.

It is acknowledged that the Bayesian approach is not without its own limitations. Historically, non-trivial problems were not solvable analytically due to the high dimensional integrals required. However, with the relatively recent availability of more computational power, a number of algorithms and approaches are now available that allow these problems to be approximated. This has seen increased uptake of Bayesian approaches within the fields of systems biology and genetics, where the inherently complex models and noisy data that these fields involve are particularly well suited to being analysed through the Bayesian approach. As long as a statistical model can be used to relate the relevant quantities, Bayesian inference can be used to give full probabilistic information on all unobserved model variables.

One of the main drawbacks to this method is that the number of model runs required to have sufficient samples in the posterior may be prohibitively high, especially where the tolerance is low or the prior distribution is very different from the posterior distribution.

This requirement for a large number of simulations for a reliable posterior is seen in all of the datasets used here. For the simulated ‘healthy’ data, the model was run 10,000,000 times in order to achieve the obtained fit. In contrast, to fit the ‘impaired’ simulated data the model was run 30,000,000 times and for the same acceptance rate the accepted particles had generally higher *ε*_NRMSE_ values. Finally, the experimental data was only able to obtain a good posterior after being run 20,000,000 times and all *ε*_NRMSE_ values were significantly above those seen in the simulated datasets. This is clearly visible in figure 11, where the distribution of *ε*_NRMSE_ values for each posterior are clearly very different. This figure clearly highlights the variance in both the error values that define a ‘good’ fit and the number of particles required for a reliable posterior distribution for different datasets.

**Fig 11.**
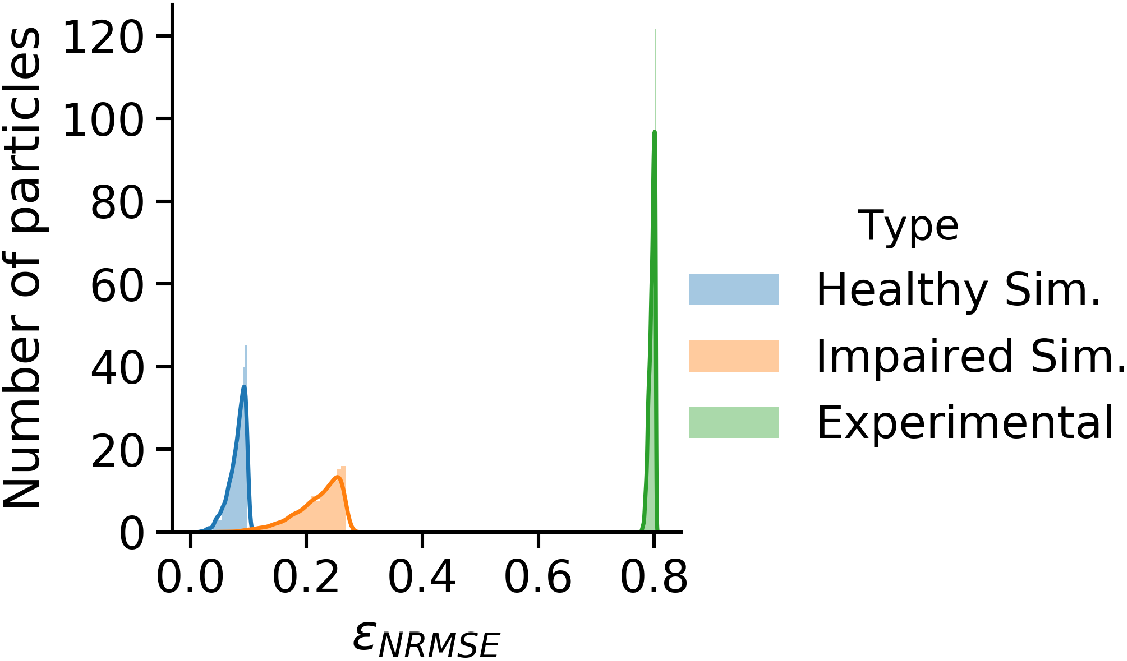
Distribution of *ε*_NRMSE_ values for the posteriors of each dataset. It can be seen here that the three datasets had very different distributions *ε*_NRMSE_ values for the particles that made up their respective posteriors. Despite this, the posterior predictive distributions for all datasets were good fits.

It should be noted that all of the obtained posterior distributions produce what are considered good fits, with those obtained for the simulated datasets far more accurate than we would ever expect to achieve when fitting experimental data. When looking at the experimental data in particular, despite the *ε*_NRMSE_ values being much higher than in the simulated data, the obtained fit captures all important behaviour and phenomena, with three clear hypoxia events visible in the inferred data trace.

More efficient methods of ABC, may alleviate the problem of requiring so many model runs to obtain good posteriors. An approach based on MCMC is more efficient than ABC REJ but the chain may become stuck in regions of low probability for long periods of time [40]. In order to deal with this problem and also the disadvantages of the rejection algorithm, an approach based on sequential Monte Carlo (ABC SMC) [37] was first proposed by Sisson et al. [41], as well as Beaumont et al. [42] and Cappé et al. [43]. In this approach, a number of sampled parameter values, known as *particles*, are sampled from the prior distribution and then propagated through a number of intermediate distributions before reaching a final target distribution. The tolerance for each successive distribution is smaller than the previous, allowing them to evolve towards the target posterior. Additionally, for a sufficiently large number of particles, the problem in MCMC of getting stuck in areas of low probability can be avoided. Developing the BayesCMD framework to use an ABC SMC approach is a key focus for future work.

## Conclusion

We have outlined how this new Bayesian framework for model analysis can be used with models of brain haemodynamics to extract information from physiological data. A more comprehensive picture of the parameter space is obtained, allowing physiological conclusions to be based on a broader picture. This is most clearly seen in the experimental data, where point estimates suggested that the values for a number of parameters had changed significantly during fitting, whilst the Bayesian method showed that the parameters were defined by a broad, roughly uniform distribution. We have also shown, through the use of data simulated from the BrainSignals model in healthy and impaired states, how the Bayesian approach allows us to better distinguish different parameter spaces. Finally, whilst we have focussed on using the BrainSignals model here, any model that can be written in a format compatible with BCMD can use this method to estimate model parameters.

A major interest within our research group is to use these models and approaches to understand and investigate further our novel measures of brain tissue physiology and metabolism and how they are linked to brain injury [44, 45]. In particular, we are interested in neonatal hypoxic ischaemic injury. The Bayesian approach provides a better representation of the parameter space and can inform a better distinction between different brain states, such as between a mild and severe injury. The method will also be adapted to use more efficient methods of parameter estimation, such as ABC SMC, reducing the number of model runs required to obtain a given tolerance.

## Supporting information

**Table 4.** Table of posterior and prior distribution information for healthy simulated data.

| Parameter | Posterior |  |  |  | Prior |  |
| --- | --- | --- | --- | --- | --- | --- |
|  | Median | Lower Quartile | Upper Quartile | IQR | Prior Min. | Prior Max. |
| sigma_coll | 65.04 | 49.4 | 79.87 | 30.47 | 31.395 | 94.185 |
| R_auto | 1.353 | 1.053 | 1.721 | 0.6679 | 0.75 | 2.25 |
| n_h | 2.541 | 1.988 | 3.079 | 1.09 | 1.25 | 3.75 |
| r_t | 0.01822 | 0.01692 | 0.01938 | 0.002461 | 0.009 | 0.027 |
| mu_max | 1.014 | 0.7679 | 1.26 | 0.4922 | 0.5 | 1.5 |
| n_m | 1.992 | 1.493 | 2.369 | 0.876 | 0.915 | 2.745 |
| r_m | 0.02879 | 0.02568 | 0.03273 | 0.007059 | 0.0135 | 0.0405 |
| P_v | 4.112 | 3.095 | 5.067 | 1.972 | 2 | 6 |
| phi | 0.03609 | 0.03474 | 0.03775 | 0.003013 | 0.018 | 0.054 |
| Xtot | 8.898 | 8.463 | 9.406 | 0.943 | 4.55 | 13.65 |

**Table 5.** Table of posterior and prior distribution information for impaired simulated data.

|  | Posterior |  |  |  | Prior |  |
| --- | --- | --- | --- | --- | --- | --- |
| Parameter | Median | Lower Quartile | Upper Quartile | IQR | Prior Min. | Prior Max. |
| sigma_coll | 64.58 | 48.86 | 79.79 | 30.93 | 31.395 | 94.185 |
| R_auto | 1.488 | 1.183 | 1.831 | 0.6479 | 0.75 | 2.25 |
| n_h | 2.337 | 1.742 | 3.014 | 1.272 | 1.25 | 3.75 |
| r_t | 0.0136 | 0.01183 | 0.01508 | 0.00325 | 0.009 | 0.027 |
| mu_max | 1.076 | 0.8283 | 1.289 | 0.4609 | 0.5 | 1.5 |
| n_m | 1.928 | 1.471 | 2.344 | 0.873 | 0.915 | 2.745 |
| r_m | 0.02738 | 0.02372 | 0.03174 | 0.00802 | 0.0135 | 0.0405 |
| P_v | 4.022 | 2.966 | 4.964 | 1.998 | 2 | 6 |
| phi | 0.03722 | 0.03349 | 0.04122 | 0.007732 | 0.018 | 0.054 |
| Xtot | 8.764 | 7.534 | 10.18 | 2.646 | 4.55 | 13.65 |

**Table 6.** Table of posterior and prior distribution information for experimental data.

| Parameter | Posterior |  |  |  | Prior |  |
| --- | --- | --- | --- | --- | --- | --- |
|  | Median | Lower Quartile | Upper Quartile | IQR | Prior Min. | Prior Max. |
| n_m | 2.078 | 1.645 | 2.446 | 0.8011 | 0.915 | 2.745 |
| r_m | 0.02574 | 0.02342 | 0.02841 | 0.004988 | 0.0135 | 0.0405 |
| K_sigma | 8.195 | 6.701 | 9.663 | 2.961 | 5 | 15 |
| p_tot | 18.48 | 17.45 | 19.4 | 1.943 | 10 | 30 |
| k_aut | 0.9799 | 0.9073 | 1.074 | 0.167 | 0.5 | 1.5 |
| v_cn | 47.88 | 43.93 | 52.13 | 8.199 | 20 | 60 |
| sigma_e0 | 0.1386 | 0.1043 | 0.1739 | 0.06966 | 0.07125 | 0.2137 |
| k2_n | 3870 | 2903 | 4822 | 1919 | 1957.83 | 5873.52 |
| Xtot | 7.067 | 6.589 | 7.555 | 0.9663 | 4.55 | 13.65 |
| R_autc | 1.955 | 1.516 | 2.517 | 1.001 | 1.1 | 3.3 |

## Acknowledgments

I.T. and J.R-B. are funded by Wellcome Trust (104580/Z/14/Z). C.P.B. is supported by a Wellcome Trust Senior Research Fellowship (209409/Z/17/Z)

